# Automated segmentation of epilepsy surgical resection cavities: comparison of four methods to manual segmentation

**DOI:** 10.1101/2024.05.13.593855

**Authors:** Merran R. Courtney, Benjamin Sinclair, Andrew Neal, John-Paul Nicolo, Patrick Kwan, Meng Law, Terence J. O’Brien, Lucy Vivash

**Affiliations:** Department of Neuroscience, School of Translational Medicine, Monash University, Melbourne, Victoria, Australia; Department of Neurology, Alfred Health, Melbourne, Victoria, Australia; Department of Neurology, Royal Melbourne Hospital, Melbourne, Victoria, Australia; Department of Medicine, The University of Melbourne, Melbourne, Victoria, Australia; Department of Radiology, Alfred Health, Melbourne, Victoria, Australia; Department of Electrical and Computer Systems Engineering, Monash University, Melbourne, Victoria, Australia

**Author notes:** **Corresponding author:** Dr Lucy Vivash, Department of Neuroscience, School of Translational Medicine, Monash University, 99 Commercial Road, Melbourne, 3004, Australia. These authors contributed equally (Merran R. Courtney, Benjamin Sinclair).

**Keywords:** Epilepsy, Automated segmentation, MRI, Neurosurgery, Resection cavity

## Abstract

Accurate resection cavity segmentation on MRI is important for neuroimaging research involving epilepsy surgical outcomes. Manual segmentation, the gold standard, is highly labour intensive. Automated pipelines are an efficient potential solution; however, most have been developed for use following temporal epilepsy surgery. Our aim was to compare the accuracy of four automated segmentation pipelines following surgical resection in a mixed cohort of subjects following temporal or extra temporal epilepsy surgery. We identified 4 open-source automated segmentation pipelines. Epic-CHOP and ResectVol utilise SPM-12 within MATLAB, while Resseg and Deep Resection utilise 3D U-net convolutional neural networks. We manually segmented the resection cavity of 50 consecutive subjects who underwent epilepsy surgery (30 temporal, 20 extratemporal). We calculated Dice similarity coefficient (DSC) for each algorithm compared to the manual segmentation. No algorithm identified all resection cavities. ResectVol (n=44, 88%) and Epic-CHOP (n=43, 86%) were able to detect more resection cavities than Resseg (n=22, 44%, P<0.001) and Deep Resection (n=21, 42%, P<0.001). The SPM-based pipelines (Epic-CHOP and ResectVol) performed better than the deep learning-based pipelines in the overall and extratemporal surgery cohorts, however there was no difference between methods in the temporal surgery cohort. These pipelines could be applied to machine learning studies of outcome prediction to improve efficiency in pre-processing data, however human quality control is still required.

## 1. Introduction

Epilepsy surgery is the most effective treatment choice for selected patients with drug resistant focal epilepsy, as it offers a better chance of long-term seizure control compared to best medical therapy. In their Cochrane systematic review, West et al found that 64% patients who underwent resective epilepsy surgery became seizure free (West et al., 2019). In order to further improve the proportion of patients for whom epilepsy surgery is successful, it is imperative that new tools are developed to improve patient selection and/or assist with tailoring a more individualised resection to maximise the chance of seizure freedom without negatively impacting neurocognitive function.

Some neuroimaging factors, including the detection of a resectable lesion on preoperative MRI, or the detection of localising ^18^F-FDG-PET hypometabolism, are associated with an increased chance of seizure freedom following epilepsy surgery (West et al., 2019, Alim-Marvasti et al., 2022, Tonini et al., 2004, Tellez-Zenteno et al., 2010, Courtney et al., 2024). Therefore, the development of neuroimaging based predictive tools may help to better delineate the subset patients who would benefit from epilepsy surgery. Accurate segmentation of the epilepsy surgery resection cavity is essential in order to incorporate data about the resection cavity into such a tool. Furthermore, surgical factors, such as complete resection of certain epileptogenic lesions, for example, focal cortical dysplasias (Rowland et al., 2012) and low grade epilepsy associated tumours (Shan et al., 2018), as well as complete resection of the epileptogenic zone (West et al., 2019, Krucoff et al., 2017), are associated with better outcome following epilepsy surgery. Accurate resection cavity segmentation is also important in assessing these factors in epilepsy surgery research. Manual segmentation is the gold standard for brain region of interest segmentation, however it is labour intensive, which prohibits its use in larger datasets, such as those required for the development of deep learning-based tools. Several open-source automated tools for segmenting the epilepsy surgery resection cavity are available, however these have largely been developed and/or validated for use following temporal epilepsy surgery (Cahill et al., 2019, Casseb et al., 2021, Arnold et al., 2022, Pérez-García et al., 2021).

The aim of the study reported here was to assess the accuracy of publicly available automated epilepsy surgery resection cavity segmentation tools compared to manual segmentation across a mixed cohort of subjects following epilepsy surgery, including subjects who have had temporal or extratemporal epilepsy surgery.

## 2. Methods

### 2.1 Ethics approval

This study was approved by the Human Research Ethics Committees (HRECs) at the Alfred Hospital (central reference number: HREC/88862/Alfred-2022-335736, local reference number: 507/22) and Royal Melbourne Hospital (local reference number: 2022.355). The HRECs determined that participant consent was not required for this study, as it utilised retrospective data acquired for clinical purposes.

### 2.2 Subject selection

We identified subjects from the Comprehensive Epilepsy Programmes at the Alfred and Royal Melbourne Hospitals in Melbourne, Australia who underwent resective epilepsy surgery for drug resistant focal epilepsy. The inclusion criteria were: (1) ≥ 16 years old at the time of surgery, (2) a preoperative T1-weighted MRI had been performed, and (3) a post-operative T1-weighted MRI had been performed ≥ 3 months following resective epilepsy surgery. For subjects meeting the inclusion criteria, we consecutively selected 30 who underwent resective temporal epilepsy surgery between January 2019 and May 2023, and 20 who underwent resective extratemporal epilepsy surgery between July 2015 and May 2023. 16 subjects who had temporal epilepsy surgery at our institutions during this period were excluded as there was no postoperative MRI performed ≥ 3 months after surgery. No subjects with extratemporal surgical resections were excluded.

### 2.3 Clinical data

Clinical data was collected from medical records. We collected demographics (age and sex), MRI diagnosis, resection type, and histopathological diagnosis. Resection type was subcategorised into anterior temporal lobectomy with amygdalohippocampectomy, temporal polectomy with encephalocoele(s) disconnection, lesionectomy, and corticectomy. Lesionectomy was defined as a sub-lobar resection that included resection of a lesion that had been detected on MRI, which may also have included resection of part of the cortex. Corticectomy was defined as a sub-lobar resection of cerebral cortex for subjects in whom the MRI did not detect a resectable lesion. The temporal polectomy with encephalocoele(s) disconnection subgroup was differentiated from the lesionectomy and corticectomy subgroups because an encephalocoele is an area of brain herniation into a skull defect, rather than a lesion within the cortex or subcortical structures, and as such we wanted to explore how each algorithm performed with an abnormality of the brain surface structure.

### 2.4 MRI acquisition

Preoperative and postoperative MRI scans were obtained as part of routine clinical care. MRI scans at the Alfred Hospital were performed on a Magnetom Skyra 3 T (n=37), while MRI scans at the Royal Melbourne Hospital were performed on either a Magnetom Skyra 3 T (n=8), Magnetom Sola 1.5 T (n=2) or Magnetom Trio 3 T (n=4; Siemens Medical Solutions, Erlangen, Germany). Three-dimensional, T1-weighted, magnetisation-prepared rapid acquisition gradient echo (MPRAGE) sequences were used for image processing. Voxel size for images performed on both Skyras and the Sola were 1.0mm^3^, while the voxel size for images performed on the Trio were 0.45 x 0.9 x 0.45 mm.

### 2.5 Manual segmentation

The resection cavity for each postoperative T1-weighted MRI was manually segmented using ITK-SNAP (Yushkevich et al., 2006) by a neurologist with expertise in epilepsy (MRC). The first five manual segmentations were checked by a senior neurologist with expertise in epilepsy (AN).

### 2.6 Automated segmentation

We performed a literature search using Medline and Web of Science databases in August 2023, to identify publicly available automated epilepsy surgery resection cavity segmentation tools. The search terms included variations and expansions on the following terms: “neuroimaging,” “magnetic resonance imaging,” “segmentation,” “epilepsy,” and “neurosurgery.” Four tools were identified. These included (1) Epilepsy Cavity Characterisation Pipeline (Epic-CHOP) (Cahill et al., 2019), (2) ResectVol (Casseb et al., 2021), (3) Deep Resection (Arnold et al., 2022), and (4) Resseg (Pérez-García et al., 2021). Epic-CHOP and ResectVol are both automated pipelines for resection cavity segmentation using Statistical Parametric Mapping (SPM) software, version 12 (Friston et al., 1994). SPM12 was used within MATLAB R2021a (Natuck, Massachusetts, USA). Deep Resection and Resseg are both deep learning models that utilise U-net convolutional neural networks to generate a segmented mask of the resection cavity. The processing pipelines for each method have been reported on in detail previously (Cahill et al., 2019, Arnold et al., 2022, Pérez-García et al., 2021, Casseb et al., 2021).

#### 2.6.1 Epic-CHOP

Using the Epic-CHOP pipeline (Cahill et al., 2019), the resection cavity was generated by deriving the difference of the preoperative and postoperative images. Briefly, the steps included: (1) recentering using the centre of mass, (2) non-linear registration of preoperative and postoperative images using SPM’s longitudinal registration toolbox, (3) segmentation into grey matter, white matter, and cerebrospinal fluid (CSF), (4) skull stripping, (5) thresholding and subtraction of the combined grey and white matter partitions of the preoperative and postoperative images, (6) erosion of the difference map by one voxel, (7) selection of three largest clusters, and (8) dilation by two voxels. Steps 6 and 8 help to reduce the segmentation and registration error in the difference map. In the automated version of Epic-CHOP, the largest cluster of the difference map is then selected as the mask of the resection cavity. However, Epic-CHOP also has the capacity to save multiple different clusters as separate segmentation masks and can therefore be used in a semi-automated way, where the user reviews each cluster and determines the cluster which best represents the resection cavity.

The Epic-CHOP algorithm generates the mask of the resection cavity in preoperative space. However, outputs in postoperative space and Montreal Neurological Institute (MNI) space are also provided, the first by applying the postoperative deformation field from the longitudinal registration, and the second by registration to SPM12’s MNI atlas.

#### 2.6.2 ResectVol

Using the ResectVol algorithm (Casseb et al., 2021), the mask of the resection region was also generated by deriving the difference between the preoperative and postoperative MR images. Briefly, the steps included: (1) linear registration to co-register the preoperative and postoperative MR images, (2) segmentation, (3) thresholding of the grey matter and white matter partitions, (4) binarization, and (5) subtraction of the preoperative and postoperative images from each other to generate the mask of the resection region. The ResectVol algorithm generates the mask of the resection cavity in preoperative space, however, an output in MNI space is also provided by registration to SPM12’s MNI atlas.

ResectVol does not have an output in postoperative space. Therefore, in order to compare the output of each algorithm to a manual segmentation performed in postoperative space, we applied the transformation matrix generated by the linear registration step in ResectVol’s algorithm to the mask of the resection cavity to move it to postoperative space.

#### 2.6.3 Deep Resection and Resseg

Deep Resection (Arnold et al., 2022) and Resseg (Pérez-García et al., 2021) are deep learning-based automated segmentation tools, which utilise three-dimensional U-net convolutional neural networks to generate a mask of the resection cavity. Deep Resection was developed using a small dataset (n=45) of subjects who underwent resective temporal epilepsy surgery and fine-tuned using a resective extratemporal epilepsy surgery dataset (n=16). Resseg was trained using the EPISURG (Pérez-García et al., 2020) database (n=430), which includes subjects who have undergone either resective temporal or extratemporal epilepsy surgery. The output mask of the resection cavity for both algorithms is generated in postoperative space. Neither algorithm requires a preoperative MRI.

#### 2.6.4 Source code availability

The source code for each algorithm is available online:

- Epic-CHOP: https://github.com/iBrain-Lab/EPIC-CHOP
- ResectVol: https://www.lniunicamp.com/resectvol
- Deep Resection: https://github.com/penn-cnt/DeepResection
- Resseg: https://github.com/fepegar/resseg

### 2.7 Data analysis

For the primary analysis we calculated the Dice similarity coefficient (DSC) for each automated segmentation mask compared to the manual segmentation in postoperative space. Secondary analyses included: (1) a comparison of the automated segmentation mask to manual segmentation in postoperative space for temporal and extratemporal subgroups, (2) a comparison of the automated segmentation mask to manual segmentation in postoperative space for different resection type subgroups, and (3) a comparison of the automated segmentation mask to manual segmentation in preoperative and MNI spaces for the algorithms which generated images in these spaces (Epic-CHOP and ResectVol). We performed the analyses in preoperative and MNI spaces as both ResectVol and Epic-CHOP generate the mask of the resection cavity in preoperative space, and each also provide the output in MNI space. As such, we chose to assess each algorithm’s performance in its native space, and all other spaces in which it provided an output. As the manual segmentation was performed in postoperative space, it was moved to preoperative space by applying the deformation field or transformation matrix from the registration process of each algorithm to enable each algorithm to be compared to manual segmentation in preoperative space. The manual segmentation was also moved to MNI space using the given algorithm’s registration to SPM12’s MNI atlas.

Finally, we explored the relationship between the volume of resection and the DSC for each algorithm. Volume of the manual segmentation was used as a proxy measure of the volume of the resection cavity. As a supplementary analysis, we assessed the performance of Epic-CHOP’s semi-automated function. We reviewed each of the three largest clusters, and visually determined which aligned with the resection cavity, and subsequently calculated the DSC compared to manual segmentation.

### 2.8 Statistical analysis

The DSC was calculated using Python according to the following equation: 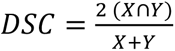, where X is the binary segmentation mask of the resection cavity generated by the automated algorithm and Y is the manual segmentation. We considered the automated algorithm to have identified the resection cavity if the DSC>0, reflecting any overlap of the two images. However, we also report the detection rates and accuracy of each algorithm where the DSC>0.5, reflecting a more meaningful overlap of at least 50%. We calculated the median DSC and interquartile range (IQR) for the total cohort and identified cohorts (DSC>0 and DSC>0.5) for the primary analysis and each of the secondary analyses. All other statistical analyses and graphs were performed using GraphPad Prism version 10.0 (Boston, Massachusetts, USA) for Mac. The detection rates (DSC>0) for each algorithm were compared using Fisher’s exact test. The distribution of data was assessed using D’Agostino-Pearson and Shapiro-Wilk normality tests. The comparison of medians between different algorithms for the primary analysis and secondary analyses with three or more groups were performed using non-parametric one-way ANOVA (Kruskal-Wallis) test, followed by pairwise comparison using Dunn’s test, with p values corrected for multiple comparisons using the Bonferroni adjustment. The comparison of medians between different algorithms for secondary analyses with two groups were performed using non-parametric independent t (Mann-Whitney) test. The relationship between volume of resection and DSC was graphically presented with a scatterplot, and a linear regression model was calculated for each algorithm.

## 3. Results

### 3.1 Population clinical characteristics

Our dataset was comprised of 50 subjects with drug resistant epilepsy who underwent temporal (n=30) or extratemporal (n=20) epilepsy surgery. The population demographic data are presented in Table 1, and individual subject characteristics are available in Appendix Table 1. The mean age at epilepsy surgery was 35.1 years. An epileptogenic lesion was detected on the preoperative MRI of 86% subjects, with the most common findings being low-grade epilepsy associated tumour (n=11) and focal cortical dysplasia (n=11), followed by cavernoma (n=7), hippocampal sclerosis (n=6), encephalocoele (n=6), and periventricular nodular heterotopia (n=2). Frontal resections were the most common extratemporal resections (n=17). A variety of resection types were represented, with the most common being lesionectomy (n=22), followed by anterior temporal lobectomy with amygdalohippocampectomy (n=18), temporal polectomy with encephalocoele(s) disconnection (n=6), and corticectomy (n=4).

**Table 1:**
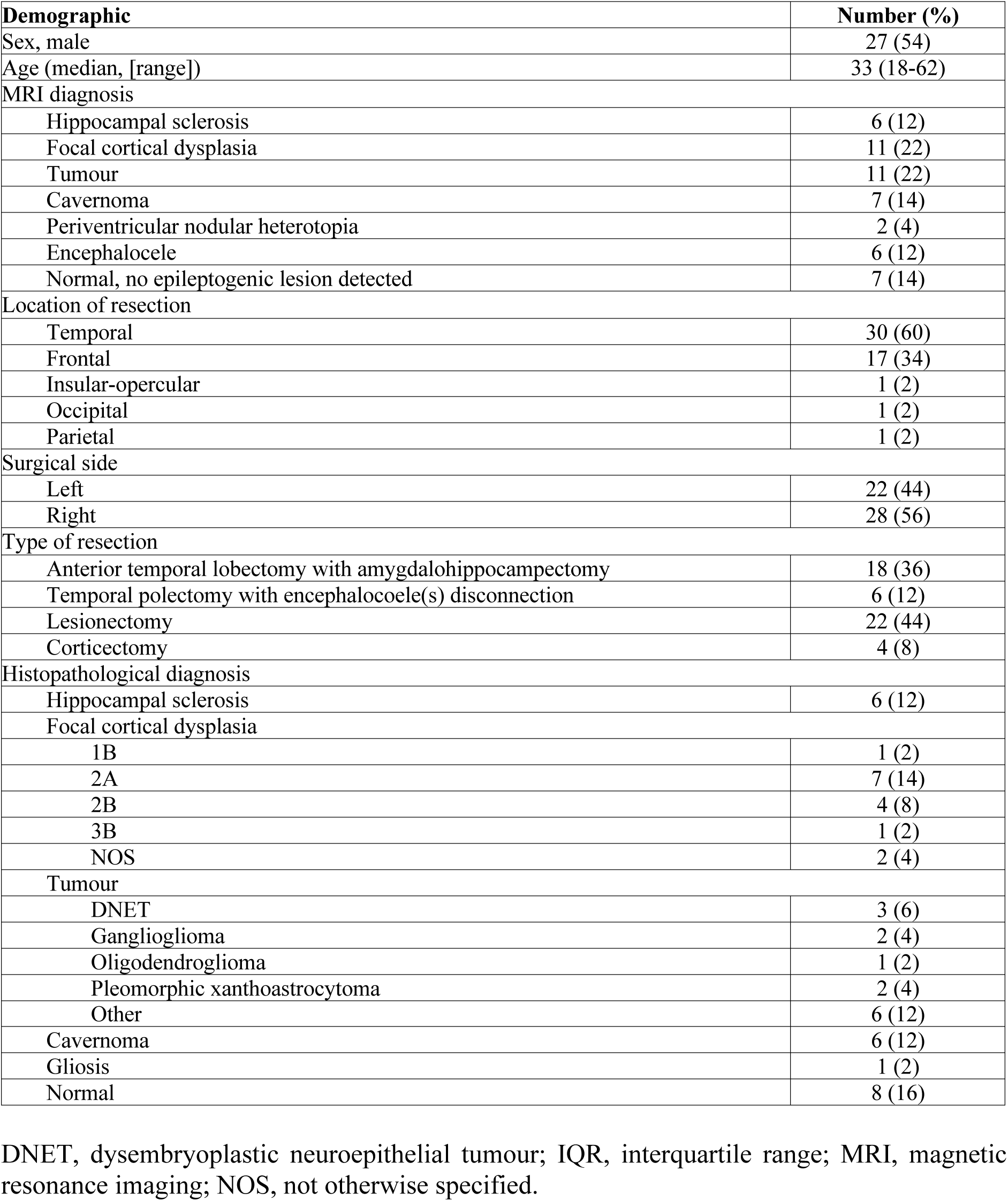
Demographic data (n=50)

### 3.2 Model performances in postoperative space

The results of the primary analysis are presented in Table 2. Example outputs from each automated segmentation algorithm are presented in Figure 1. None of the algorithms identified (DSC>0) every resection cavity in our cohort (n=50). Epic-CHOP (n=43, 86%) and ResectVol (n=44, 88%) both identified more resection cavities than either Resseg (n=22, 44%, p<0.001) or Deep Resection (n=21, 42%, p<0.001). There was a difference (p<0.001) between the DSCs (median [IQR]) of the four algorithms: Epic-CHOP 0.71 (0.24), ResectVol 0.67 (IQR, 0.34), Resseg 0.00 (IQR, 0.84), and Deep Resection 0.00 (IQR, 0.58). However, when adjusted for multiple comparisons, there was only a significant difference between Epic-CHOP and Deep Resection (p<0.05), and ResectVol and Deep Resection (p<0.05). Raw data from pairwise comparisons for all analyses are presented in Appendix Table 2.

**Figure 1:**
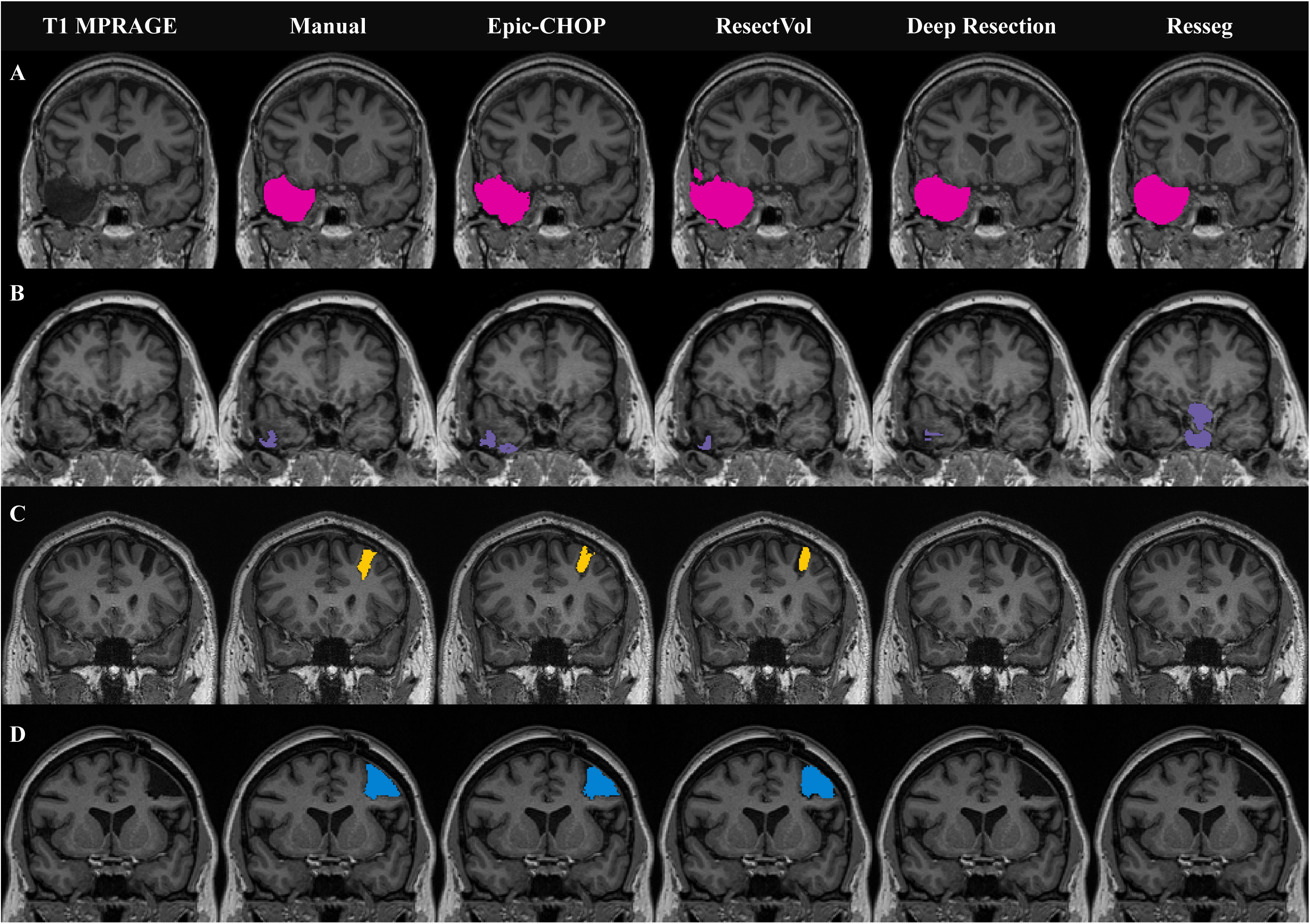
Automated segmentation outputs for each model. The automated resection cavity segmentation output overlayed on the postoperative T1-weighted MPRAGE for each algorithm for different resection types: (A) right anterior temporal lobectomy (DSC range 0.67-0.90); (B) right temporal polectomy with encephalocoele disconnection (DSC range 0.00-0.51); (C) left frontal corticectomy (0.00-0.63); (D) left frontal lesionectomy (DSC range 0.00-0.85).

**Table 2:** Dice similarity coefficient for each automated segmentation algorithm compared to manual segmentation in post-operative space for the total cohort, temporal epilepsy surgery and extratemporal epilepsy surgery groups.

| Algorithm | Total cohort |  |  |  |  | Temporal epilepsy surgery |  |  |  |  | Extratemporal epilepsy surgery |  |  |  |  |
| --- | --- | --- | --- | --- | --- | --- | --- | --- | --- | --- | --- | --- | --- | --- | --- |
|  | No. subjects identified |  | Dice similarity coefficient (median [IQR]) |  |  | No. subjects identified |  | Dice similarity coefficient (median [IQR]) |  |  | No. subjects identified |  | Dice similarity coefficient (median [IQR]) |  |  |
|  | DSC >0 | DSC >0.5 | Total cohort | Cohort with DSC>0 | Cohort with DSC>0.5 | DSC >0 | DSC >0.5 | Total cohort | Cohort with DSC>0 | Cohort with DSC>0.5 | DSC >0 | DSC >0.5 | Total cohort | Cohort with DSC>0 | Cohort with DSC>0.5 |
| Epic-CHOP | 43/50 | 38/50 | 0.71<br>(0.24) | 0.74<br>(0.17) | 0.77<br>(0.13) | 28/30 | 26/30 | 0.77<br>(0.21) | 0.78<br>(0.16) | 0.77<br>(0.21) | 15/20 | 12/20 | 0.63<br>(0.74) | 0.71<br>(0.16) | 0.73<br>(0.11) |
| ResectVol | 44/50 | 36/50 | 0.67<br>(0.34) | 0.72<br>(0.23) | 0.67<br>(0.34) | 27/30 | 22/30 | 0.67<br>(0.29) | 0.68<br>(0.23) | 0.68<br>(0.22) | 17/20 | 14/20 | 0.70<br>(0.60) | 0.72<br>(0.20) | 0.75<br>(0.13) |
| Resseg | 22/50 | 22/50 | 0.00<br>(0.84) | 0.85<br>(0.08) | 0.85<br>(0.08) | 19/30 | 19/30 | 0.81<br>(0.87) | 0.86<br>(0.07) | 0.86<br>(0.07) | 3/20 | 3/20 | 0.00<br>(0.00) | 0.76<br>(0.14) | 0.76<br>(0.14) |
| Deep Resection | 21/50 | 16/50 | 0.00<br>(0.58) | 0.68<br>(0.38) | 0.75<br>(0.25) | 18/30 | 14/30 | 0.40<br>(0.74) | 0.62<br>(0.44) | 0.81<br>(0.29) | 3/20 | 2/20 | 0.00<br>(0.00) | 0.68<br>(0.15) | 0.71<br>(0.03) |

DSC, Dice similarity coefficient; IQR, interquartile range.

### 3.3 Model performances for temporal and extratemporal resection subgroups

The results of the temporal (n=30) and extratemporal (n=20) resection subgroup analyses are presented in Table 2. Epic-CHOP (n=28, 93%) and ResectVol (n=27, 90%) identified (DSC>0) more temporal resection cavities than either Resseg (n=19, 63%, p<0.05) or Deep Resection (n=18, 60%, p<0.05). There was no difference (p=0.13) between the DSCs (median [IQR]) for the temporal epilepsy surgery cohort: Epic-CHOP 0.77 (0.21), ResectVol 0.67 (0.29), Resseg 0.81 (0.87), and Deep Resection 0.40 (0.74).

For the extratemporal epilepsy surgery subgroup, Epic-CHOP (n=15, 75%) and ResectVol (n=17, 85%) identified (DSC>0) more resection cavities than either Resseg (n=3, 15%, p<0.001) or Deep Resection (n=3, 15%, p<0.001). There was a difference (p<0.001) in the DSCs (median [IQR]) for the extratemporal epilepsy surgery cohort: Epic-CHOP 0.63 (0.74), ResectVol 0.70 (0.60), Resseg 0.00 (0.00) and Deep Resection 0.00 (0.00). The results of pairwise comparisons are presented in Appendix Table 2.

### 3.4 Model performances in postoperative space for different resection types

The results of the subgroup analyses for different resection types are presented in Table 3. There was a difference in the median DSCs for each algorithm for the anterior temporal lobectomy with amygdalohippocampectomy (n=18, p=0.01), lesionectomy (n=22, p=0.01) and corticectomy (n=4, p=0.03) subgroups. However, pairwise comparisons did not demonstrate any significant differences between algorithms following adjustment for multiple comparisons (Appendix Table 3). There was no difference in the median DSCs for each algorithm in the temporal polectomy with encephalocoele(s) disconnection (n=6) subgroup.

**Table 3:** Dice similarity coefficient for each automated segmentation algorithm compared to manual segmentation in post-operative space for each resection type.

| Type of resection | Algorithm | No. subjects identified |  | Dice similarity coefficient (median [IQR]) |  |  |
| --- | --- | --- | --- | --- | --- | --- |
|  |  | DSC>0 | DSC>0.5 | Total cohort | Cohort with DSC>0 | Cohort with DSC>0.5 |
| ATL+AH | Epic-CHOP | 18/18 | 17/18 | 0.79 (0.11) | 0.79 (0.11) | 0.79 (0.10) |
|  | ResectVol | 17/18 | 15/18 | 0.68 (0.19) | 0.71 (0.17) | 0.76 (0.14) |
|  | Resseg | 15/18 | 15/18 | 0.84 (0.13) | 0.86 (0.08) | 0.86 (0.08) |
|  | Deep Resection | 12/18 | 9/18 | 0.51 (0.80) | 0.68 (0.37) | 0.72 (0.32) |
| Temporal polectomy + encephalocele disconnection | Epic-CHOP | 4/6 | 3/6 | 0.48 (0.45) | 0.55 (0.05) | 0.56 (0.04) |
|  | ResectVol | 4/6 | 2/6 | 0.40 (0.44) | 0.50 (0.21) | 0.67 (0.10) |
|  | Resseg | 0/6 | 0/6 | 0.00 (0.00) | - | - |
|  | Deep Resection | 2/6 | 1/6 | 0.04 (0.20) | 0.42 (0.17) | 0.59 (0.00) |
| Lesionectomy | Epic-CHOP | 18/22 | 16/22 | 0.68 (0.67) | 0.77 (0.16) | 0.78 (0.16) |
|  | ResectVol | 20/22 | 16/22 | 0.73 (0.27) | 0.74 (0.18) | 0.76 (0.11) |
|  | Resseg | 7/22 | 7/22 | 0.00 (0.72) | 0.84 (0.07) | 0.84 (0.07) |
|  | Deep Resection | 8/22 | 6/22 | 0.00 (0.63) | 0.75 (0.24) | 0.81 (0.14) |
| Corticectomy | Epic-CHOP | 3/4 | 3/4 | 0.70 (0.19) | 0.71 (0.02) | 0.71 (0.02) |
|  | ResectVol | 3/4 | 3/4 | 0.64 (0.24) | 0.67 (0.10) | 0.64 (0.10) |
|  | Resseg | 0/4 | 0/4 | 0.00 (0.00) | - | - |
|  | Deep Resection | 0/4 | 0/4 | 0.00 (0.00) | - | - |

ATL+AH, anterior temporal lobectomy with amygdalohippocampectomy; DSC, Dice similarity coefficient; IQR, interquartile range.

### 3.5 Model performances in preoperative and MNI spaces

The model performances for Epic-CHOP and ResectVol in preoperative and MNI space are presented in Table 4. There was no difference between the DSCs (median [IQR]) in preoperative space (p=0.56): Epic-CHOP 0.71 (0.28), ResectVol 0.68 (0.35) nor MNI space (p=0.86): Epic-CHOP 0.71 (0.27), ResectVol 0.67 (0.34).

**Table 4:**
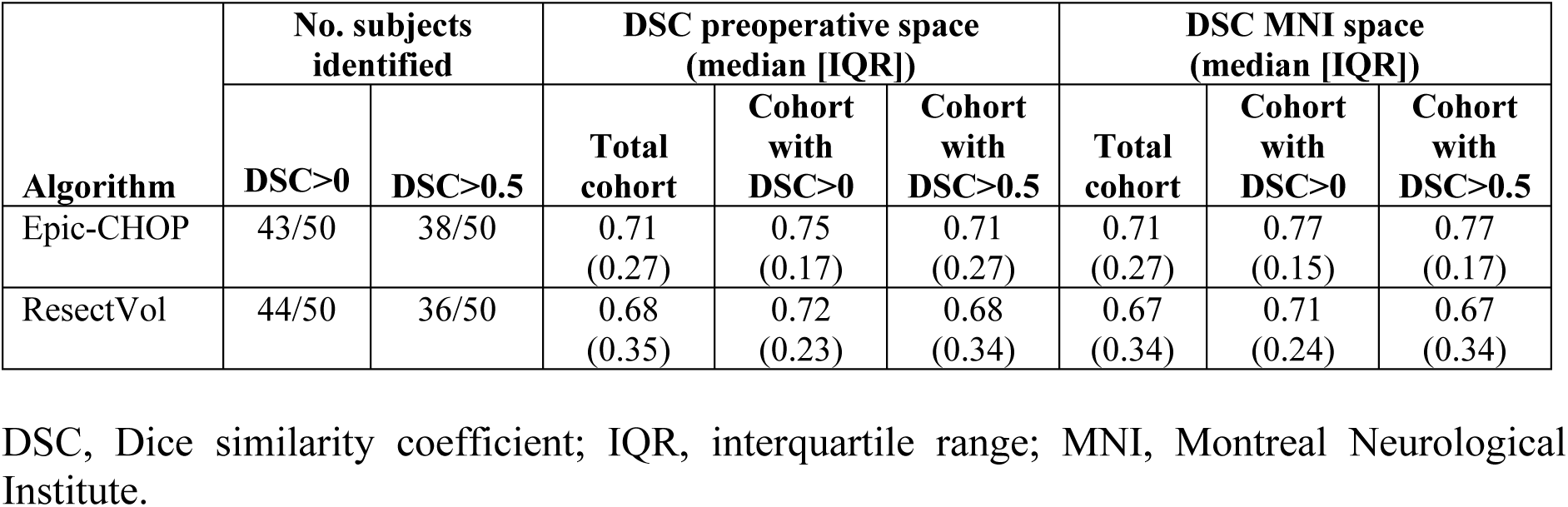
Dice similarity coefficient for automated segmentation outputs in preoperative space and MNI space compared to manual segmentation.

| Algorithm | No. subjects identified |  | DSC preoperative space (median [IQR]) |  |  | DSC MNI space (median [IQR]) |  |  |
| --- | --- | --- | --- | --- | --- | --- | --- | --- |
|  | DSC>0 | DSC>0.5 | Total cohort | Cohort with DSC>0 | Cohort with DSC>0.5 | Total cohort | Cohort with DSC>0 | Cohort with DSC>0.5 |
| Epic-CHOP | 43/50 | 38/50 | 0.71 (0.27) | 0.75 (0.17) | 0.71 (0.27) | 0.71 (0.27) | 0.77 (0.15) | 0.77 (0.17) |
| ResectVol | 44/50 | 36/50 | 0.68 (0.35) | 0.72 (0.23) | 0.68 (0.34) | 0.67 (0.34) | 0.71 (0.24) | 0.67 (0.34) |

DSC, Dice similarity coefficient; IQR, interquartile range; MNI, Montreal Neurological Institute.

### 3.6 Relationship between volume of resection and DSC

We examined the relationship between volume of resection and DSC for each algorithm in scatterplots, presented in Figure 2. Linear regression analyses demonstrated a weak relationship for all algorithms, such that DSC increased with increasing resection volume: Epic-CHOP (R^2^ = 0.24, p<0.001), ResectVol (R^2^ = 0.12, p=0.02), Resseg (R^2^ = 0.24, p<0.001), and Deep Resection (R^2^ = 0.13, p=0.01).

**Figure 2:**
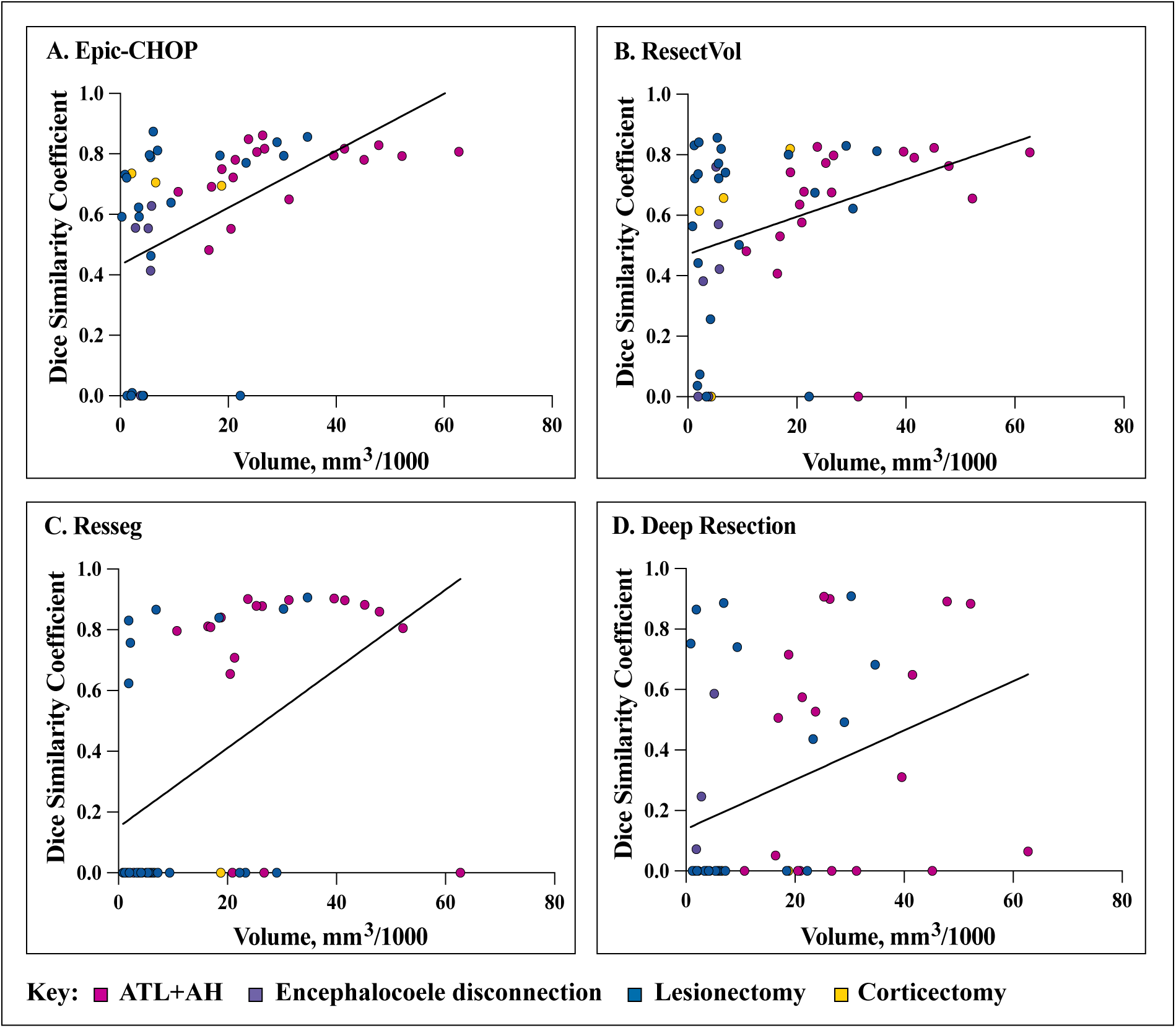
Relationship of model performance with volume of the resection cavity. Relationship between the volume of the resection cavity and Dice similarity coefficient for each algorithm with associated linear regression model: (A) Epic-CHOP, coefficient of determination (R^2^) = 0.24 (p<0.001); (B) ResectVol, R^2^ = 0.12 (p=0.02); (C) Resseg, R^2^ = 0.24 (p<0.001); (D) Deep Resection, R^2^ = 0.13 (p=0.01).

### 3.7 Analysis of Epic-CHOP semi-automated

The results of the supplementary analysis of the semi-automated version of Epic-CHOP are presented in Appendix Table 4. One additional resection region was detected compared to Epic-CHOP automated (n=44/50), however this was not statistically significant (p=0.99). There was no difference in the median DSCs between the automated and semi-automated versions of Epic-CHOP (P=1.00).

## 4. Discussion

In this study, we compare four publicly available automated tools for segmenting the epilepsy surgery resection cavity. Our results demonstrate: (1) the SPM-based tools (Epic-CHOP and ResectVol) identified more resection cavities than the deep learning-based tools (Resseg and Deep Resection) for the mixed temporal and extratemporal epilepsy surgery cohort, and had higher median DSCs than Deep Resection; (2) all four tools analysed had similar detection rates and accuracies for temporal epilepsy surgery resection cavities; (3) the SPM-based tools had higher detection rates and better accuracies than the deep learning tools in the extratemporal epilepsy surgery cohort; (4) the accuracy of each model improves as the size of the resection cavity increases; and (5) quality control is required when using all tools.

Overall, the SPM-based automated segmentation tools (Epic-CHOP and ResectVol) had higher detection rates and better accuracies than the deep learning-based automated segmentation tools (Resseg and Deep Resection) on the mixed cohort of temporal and extratemporal epilepsy surgical resections. Although there was no difference between the performance of the models in the temporal epilepsy surgery cohort, the SPM-based tools performed better than the deep learning-based tools in the extratemporal cohort. Deep Resection was primarily trained on temporal epilepsy resections, and Resseg was trained on the EPISURG dataset, which is predominantly (81%) temporal epilepsy resections, which may explain why the deep learning-based methods performed less well on the extratemporal resections compared to the temporal resections. Although not statistically significant, the deep learning model Resseg had the highest median DSC in the temporal epilepsy surgery group. Therefore, a similar model trained with extratemporal resections has the potential to be highly accurate. The deep learning models also had the fastest computational times, taking only a few minutes per subject, in comparison to the SPM-based models, which took approximately 20-25 minutes per subject. However, since all methods can be configured to loop over subjects, the overall time saved to the user is considerable for all the methods tested.

Conversely, although both SPM-based tools were developed using temporal epilepsy resections, the principles underlying these pipelines are not exclusive to temporal lobe resections, which may explain why these models also performed better on the extra-temporal epilepsy resections. Both Epic-CHOP and ResectVol generate the mask of the resection cavity by deriving the difference between the preoperative and postoperative MR images. These models rely on accurate registration and segmentation of the images into grey matter, white matter, and CSF partitions. The SPM segmentation toolbox utilises Bayesian probability to determine the tissue type, with both voxel intensity and tissue probability maps included in the calculation. Certain features on the preoperative and postoperative MR images may lead to segmentation error in the SPM-based tools. For example, blood in the resection cavity may be segmented as if it was grey matter, such that following subtraction of the postoperative grey and white matter partitions from the preoperative, the resection cavity may be missed or underestimated. We attempted to avoid this by using postoperative MRIs performed ≥ 3 months following surgery. Alternatively, certain lesions on the preoperative MRI, for example a cystic lesion, which has a similar voxel intensity as CSF, may be segmented as CSF, again leading to either a missed or underestimated resection cavity. Epic-CHOP and Resect Vol differ in two main aspects. Firstly, Epic-CHOP performs an erosion and dilation of the difference mask to account for segmentation and registration error, which may account for the slightly higher DSC in the identified lesions (Table 2). Secondly, ResectVol performs a linear registration between the preoperative and postoperative MRIs, whilst Epic-CHOP performs a non-linear registration to account for post-operative collapse of tissue into the resection cavity. However, this second aspect will not be reflected in the DSCs as the manual segmentations were performed on postoperative MRIs. Furthermore, while the default output for the Epic-CHOP and ResectVol resection region segmentations is in preoperative space, which endeavours to reflect the preoperative tissue that was resected, the two deep learning methods output the resection region in postoperative space. Although the ability to use Resseg and Deep Resection in situations where only a postoperative MRI is available is a potential advantage, the interpretation of the segmented resection region when performed in postoperative space is different to when it is performed in preoperative space. If the intention is to use the deep learning-based algorithms to make inferences about the preoperative tissue that has been resected, a further registration step to the preoperative MRI would be required.

We found that there was a weak relationship between the volume of the resection cavity and the DSC for each algorithm, such that larger resections were associated with higher DSCs. In our cohort, the largest resections were anterior temporal lobectomies with amygdalohippocampectomies (see Figure 2), which may explain this relationship, as each of the models was predominantly trained and/or developed using such resections. Smaller resections, particularly the temporal polectomies with encephalocoele(s) disconnections, were overall less accurate than the anterior temporal lobectomy with amygdalohippocampectomies. It is clear, regardless of the algorithm being used, that a quality control step, to check that the automated segmentation has not missed or grossly underestimated the resection cavity, is required.

The main limitation of this study was that the manual segmentations were only performed by a single researcher. Therefore, an inter-rater reliability could not be calculated, and it is possible that some of the differences between the manual and automated segmentations may be due to error in the manual segmentation. However, we felt this was acceptable, as the purpose of the study was not validation of the tools, but to compare them in a mixed cohort of subjects following either temporal or extratemporal epilepsy surgery. Another limitation is that most of the extratemporal resections were frontal epilepsy resections (n=17/20, 85%). We consecutively selected the 20 extratemporal cases, therefore the predominance of frontal cases reflects the fact that frontal epilepsy resections were the most common extratemporal resections in our institutions over the inclusion time-period. This raises the question about whether the extratemporal epilepsy resection results are generalisable to extra-frontal resections, however, we believe that the principles underpinning the performance of each model on the extratemporal resections would apply regardless of the extratemporal location. Thirdly, all algorithms provided an output in postoperative space except for ResectVol. Therefore, in order to compare each method, we had to move the ResectVol output from preoperative space to postoperative space. We used the transformation matrix from the linear registration used in ResectVol to minimise bias, however this process may have biased against ResectVol in the primary analysis. To assess ResectVol’s performance in its native space, we also performed analyses in preoperative and MNI spaces, and these results were similar to postoperative space.

In conclusion, automated resection cavity segmentation pipelines have the potential to reduce the significant burden of time involved in manually generating a mask of the resection cavity for research purposes. In our mixed temporal and extratemporal epilepsy resection cohort, the SPM-based automated segmentation tools (Epic-CHOP and ResectVol) performed better than the deep learning-based automated segmentation tools (Resseg and Deep Resection). However, none of the pipelines identified every resection cavity, and their use should be combined with a human quality control step.

## CONTRIBUTORS

MRC wrote the first draft of the manuscript and performed the literature search. MRC, BS, AN, J-PN, PK, ML, TJO and LV conceptualised and designed the study protocol. MRC and BS acquired and processed the data. MRC, BS, AN, J-PN, PK, ML, TJO and LV interpreted the data. All authors critically revised the manuscript for intellectual content, and all authors approve the final manuscript.

