## Appendix for "Automated segmentation of epilepsy surgical resection cavities: comparison of four methods to manual segmentation"

### Appendix Table of Contents

**Appendix Table 1:** Individual subject characteristics

| Subject No. | Age at surgery, sex | MRI diagnosis | Type of resection | Histopathological diagnosis |
|---|---|---|---|---|
| 1 | 25, M | L frontal LEAT | L frontal lesionectomy | Pleomorphic xanthoastrocytoma |
| 2 | 30, M | L temporal encephalocele | L temporal polectomy and encephalocele disconnection | Normal |
| 3 | 52, F | R temporal encephalocele | R temporal polectomy and encephalocele disconnection | Normal |
| 4 | 50, M | PVNH | R ATL + AH | Hippocampal sclerosis |
| 5 | 30, M | R MTS | R ATL + AH | Normal |
| 6 | 20, M | Normal | L frontal corticectomy | FCD 2B |
| 7 | 40, M | L frontal cavernoma | L frontal lesionectomy | Cavernoma |
| 8 | 62, F | L MTS | L ATL + AH | Hippocampal sclerosis |
| 9 | 48, F | L temporal LEAT | L ATL + AH | FCD 2, NOS |
| 10 | 34, F | Normal | L frontal corticectomy | FCD 2A |
| 11 | 45, F | Normal | R ATL + AH | FCD 2A |
| 12 | 37, M | R temporal cavernoma | R temporal lesionectomy | DNET |
| 13 | 30, M | R temporal LEAT | R ATL + AH | FCD 3B |
| 14 | 55, M | Normal | L ATL + AH | Normal |
| 15 | 25, M | R frontal glioma | R frontal lesionectomy | Glioblastoma |
| 16 | 33, M | L frontal cavernoma | L frontal lesionectomy | Cavernoma |
| 17 | 26, F | L temporal cavernoma | L temporal lesionectomy | Normal |
| 18 | 29, M | L temporal LEAT | L ATL + AH | Diffuse glioneuronal tumour with oligodendroglioma like features |
| 19 | 43, M | L temporal encephalocele | L temporal polectomy and encephalocele disconnection | Gliosis |
| 20 | 30, M | R temporal encephalocele | R ATL + AH + encephalocele disconnection | Hippocampal sclerosis |
| 21 | 26, F | R insular LEAT | R insular lesionectomy | Ganglioglioma |
| 22 | 43, F | R MTS | R ATL + AH | Hippocampal sclerosis |
| 23 | 41, M | R temporal LEAT | R ATL + AH + lesionectomy | LEAT, NOS |
| 24 | 24, M | R temporal encephalocele | R temporal polectomy and encephalocele disconnection | Normal |
| 25 | 29, M | R temporal cavernoma | R temporal lesionectomy | Cavernoma |
| 26 | 66, F | R temporal encephalocele | R temporal polectomy and encephalocele disconnection | Normal |
| 27 | 31, F | R MTS | R ATL + AH | FCD 1B |
| 28 | 33, M | L temporal cavernoma | L temporal polectomy and encephalocele disconnection | Cavernoma |
| 29 | 18, F | PVNH | L ATL + AH + lesionectomy | Pleomorphic xanthoastrocytoma |
| 30 | 40, F | R temporal LEAT | R ATL + AH | Extraventricular benign choroid plexus lesion |
| 31 | 59, F | R temporal cavernoma | R ATL + AH + lesionectomy | Cavernoma |
| 32 | 19, F | L MTS | L ATL + AH | Hippocampal sclerosis |
| 33 | 23, F | L temporal LEAT | L ATL + AH + lesionectomy | DNET |
| 34 | 37, F | Normal | L ATL + AH | Sporadic cortical hamartoma |
| 35 | 24, M | Normal | L frontal corticectomy | Normal |
| 36 | 36, M | Normal | R occipital corticectomy | Meningioangiomatosis |
| 37 | 34, M | L frontal FCD | L frontal lesionectomy | FCD 2A |
| 38 | 28, F | L MTS | L ATL + AH | Hippocampal sclerosis |
| 39 | 34, F | R temporal cavernoma | R temporal lesionectomy | Cavernoma |
| 40 | 19, F | R frontal cystic lesion | R frontal lesionectomy | DNET |
| 41 | 53, F | L frontal FCD | L frontal lesionectomy | FCD 2B |
| 42 | 28, M | R temporal FCD | R temporal lesionectomy | Gangliocytoma |
| 43 | 43, F | R frontal LEAT | R frontal lesionectomy | Oligodendroglioma |
| 44 | 34, F | R frontal FCD | R frontal lesionectomy | FCD 2B |
| 45 | 24, M | R frontal FCD | R frontal lesionectomy | FCD 2A |
| 46 | 54, M | R parietal FCD | R parietal lesionectomy | FCD 2A |
| 47 | 20, M | R frontal FCD | R frontal lesionectomy | FCD 2A |
| 48 | 20, M | R frontal FCD | R frontal lesionectomy | FCD 2A |
| 49 | 38, F | L frontal FCD | L frontal lesionectomy | FCD 2B |
| 50 | 33, M | R frontal FCD | R frontal lesionectomy | FCD, NOS |

ATL + AH, anterior temporal lobectomy with amygdalohippocampectomy; DNET, dysembryoplastic neuroepithelial tumour; FCD, focal cortical dysplasia; LEAT, low grade epilepsy associated neuroepithelial tumour; MRI, magnetic resonance imaging; MTS, mesial temporal sclerosis; PVNH, periventricular nodular heterotopia.

**Appendix Table 2:** Results of pairwise comparisons performed using Dunn's multiple comparisons test.

| | Raw p-value | Adjusted p-value |
|---|---|---|
| Primary Analysis | | |
| ResectVol vs. Epic-CHOP | >0.99 | 1.00 |
| ResectVol vs. Deep Resection | 0.005 | 0.03* |
| ResectVol vs. Resseg | 0.42 | 1.00 |
| Epic-CHOP vs. Deep Resection | 0.002 | 0.01* |
| Epic-CHOP vs. Resseg | 0.27 | 1.00 |
| Deep Resection vs. Resseg | 0.77 | 1.00 |
| Temporal Resection | | |
| ResectVol vs. Epic-CHOP | >0.99 | 1.00 |
| ResectVol vs. Deep Resection | >0.99 | 1.00 |
| ResectVol vs. Resseg | >0.99 | 1.00 |
| Epic-CHOP vs. Deep Resection | 0.27 | 1.00 |
| Epic-CHOP vs. Resseg | >0.99 | 1.00 |
| Deep Resection vs. Resseg | 0.27 | 1.00 |
| Extratemporal Resection | | |
| ResectVol vs. Epic-CHOP | >0.99 | 1.00 |
| ResectVol vs. Deep Resection | 0.0001 | 0.001* |
| ResectVol vs. Resseg | 0.0003 | 0.002* |
| Epic-CHOP vs. Deep Resection | 0.0056 | 0.03* |
| Epic-CHOP vs. Resseg | 0.0118 | 0.07 |
| Deep Resection vs. Resseg | >0.99 | 1.00 |
| ATL+AH | | |
| ResectVol vs. Epic-CHOP | >0.99 | 1.00 |
| ResectVol vs. Deep Resection | >0.99 | 1.00 |
| ResectVol vs. Resseg | 0.24 | 1.00 |
| Epic-CHOP vs. Deep Resection | 0.16 | 0.94 |
| Epic-CHOP vs. Resseg | >0.99 | 1.00 |
| Deep Resection vs. Resseg | 0.01 | 0.06 |
| Temporal polectomy with encephalocoele(s) disconnection | | |
| ResectVol vs. Epic-CHOP | >0.99 | 1.00 |
| ResectVol vs. Deep Resection | >0.99 | 1.00 |
| ResectVol vs. Resseg | 0.14 | 0.84 |
| Epic-CHOP vs. Deep Resection | >0.99 | 1.00 |
| Epic-CHOP vs. Resseg | 0.14 | 0.84 |
| Deep Resection vs. Resseg | >0.99 | 1.00 |
| Lesionectomy | | |
| ResectVol vs. Epic-CHOP | >0.99 | 1.00 |
| ResectVol vs. Deep Resection | 0.05 | 0.30 |
| ResectVol vs. Resseg | 0.07 | 0.42 |
| Epic-CHOP vs. Deep Resection | 0.18 | 1.00 |
| Epic-CHOP vs. Resseg | 0.26 | 1.00 |
| Deep Resection vs. Resseg | >0.99 | 1.00 |
| Corticectomy | | |
| ResectVol vs. Epic-CHOP | >0.99 | 1.00 |
| ResectVol vs. Deep Resection | 0.33 | 1.00 |
| ResectVol vs. Resseg | 0.33 | 1.00 |
| Epic-CHOP vs. Deep Resection | 0.18 | 1.00 |
| Epic-CHOP vs. Resseg | 0.18 | 1.00 |
| Deep Resection vs. Resseg | >0.99 | 1.00 |

**Appendix Table 3:** Results of semi-automated version of Epic-CHOP

| Space | No. Subjects identified (DSC>0) | Dice similarity coefficient (median [IQR]) |
|---|---|---|
| Post-operative | 44/50 | 0.71 (0.24) |
| Pre-operative | | 0.71 (0.27) |
| MNI | | 0.66 (0.31) |

DSC, Dice similarity coefficient; IQR, interquartile range; MNI, Montreal Neurological Institute.
